# Pharyngeal timing and particle transport defects in *Caenorhabditis elegans* feeding mutants

**DOI:** 10.1101/2021.09.30.462635

**Authors:** Isaac Ravi Brenner, David M. Raizen, Christopher Fang-Yen

## Abstract

The nematode *C. elegans* uses rhythmic muscle contractions (pumps) of the pharynx, a tubular feeding organ, to filter, transport, and crush food particles. A number of feeding mutants have been identified, including those with slow pharyngeal pumping rate, weak muscle contraction, defective muscle relaxation, and defective grinding of bacteria. Many aspects of these pharyngeal behavioral defects and how they affect pharyngeal function are not well understood. For example, the behavioral deficits underlying inefficient particle transport in ‘slippery’ mutants have been unclear. Here we use high speed video microscopy to describe pharyngeal pumping behaviors and particle transport in wild-type animals and in feeding mutants. Different ‘slippery’ mutants exhibit distinct defects including weak isthmus contraction, failure to trap particles in the anterior isthmus, and abnormal timing of contraction and relaxation in pharyngeal compartments. Our results show that multiple deficits in pharyngeal timing or contraction can cause defects in particle transport.

## Introduction

The nematode *C. elegans* consumes food bacteria using movements of the pharynx, a neuromuscular feeding organ that connects to the stoma (buccal cavity) and to the intestine (Fig. 1). The pharynx displays two behaviors: pharyngeal pumping and isthmus peristalsis. Pharyngeal pumping consists of rhythmic cycles of contraction and relaxation throughout most of the pharynx. During contraction, the radially oriented muscle fibers open the pharyngeal lumen; during relaxation the lumen closes. Isthmus peristalsis is an anterior-to-posterior wave of contraction in the posterior half of the isthmus that occurs after a fraction of pumps (Avery and Horvitz, 1987). Together, these movements filter food particles from surrounding fluids, transport the particles posteriorly, and crush them before they enter the intestine.

**Figure 1.**
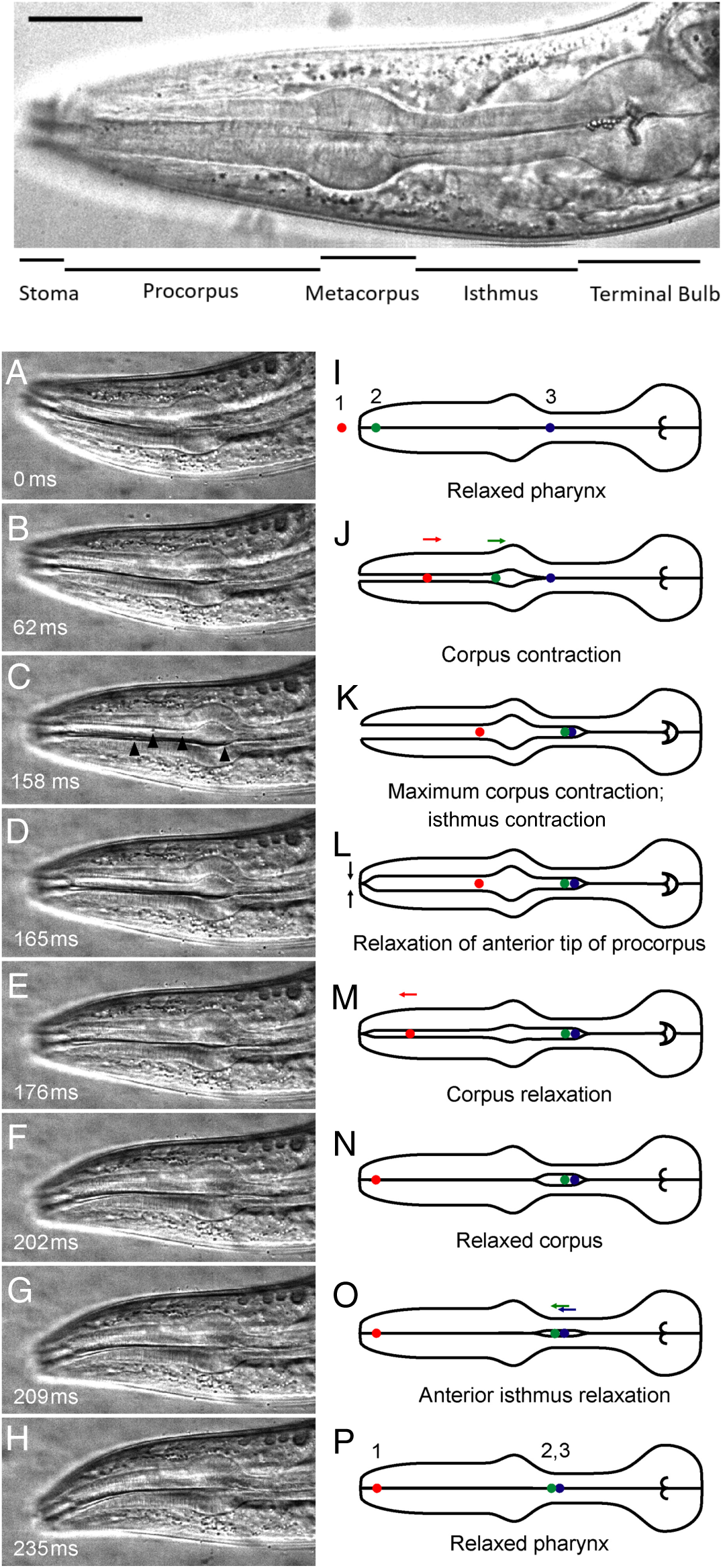
Pharyngeal anatomy and behavior. Top: Annotated DIC image of the anterior of an adult hermaphrodite (field of view 177 µm x 57 µm). The corpus includes the procorpus and metacorpus. The intestine is posterior to the terminal bulb. (A-H):Frames from high-speed video sequence showing movements of pharyngeal muscles and 0.75 µm polystyrene particles during pharyngeal pumping in adult worm. Field of view 136 µm x 65 µm. Black triangles in C indicate several particles. (I–P) Corresponding illustrations showing initial, intermediate, and final positions of three representative particles labeled 1 (red), 2 (green), and 3 (blue). During one cycle of contraction and relaxation, particle 1 has been transported from outside the pharynx to the anterior procorpus trap. Particle 2 has been transported from the anterior procorpus trap to the anterior isthmus trap. Particle 3 remains in the anterior isthmus trap. Adapted with permission from (Fang-Yen et al., 2009).

The pharynx is divided into several compartments including the anterior and posterior (terminal) bulbs. Detailed studies of pharyngeal pumping have revealed differences in timing of contractions and relaxations in these compartments (Avery, 1993a, 1993b; Fang-Yen et al., 2009). The corpus, which extends from the stoma to the anterior bulb, is the first to contract, drawing particles from the stoma into the pharyngeal lumen (Fig 1). Contraction of the anterior isthmus and then the terminal bulb follow contraction of the corpus with short delays. Pharyngeal relaxation of the various compartments proceeds in the same order as their contraction but more rapidly (Fig 1). (Fang-Yen 2009).

The sequence of relaxations in pharyngeal compartments is closely tied to the mechanism of particle trapping (Fig. 1). The anterior tip of the corpus relaxes slightly before the rest of the corpus; this allows food particles to become trapped at the anterior corpus. Similarly, relaxation of the metacorpus before the anterior isthmus allows food particles to become trapped at the anterior isthmus. At each of the two trap sites (anterior corpus and anterior isthmus), earlier relaxation of the structures immediately anterior to the trap prevents anteriorward transport of particles while fluids are expelled (Fang-Yen et al., 2009). Under some conditions such as violet light exposure, the trap at the anterior procorpus remains contracted during corpus relaxation, causing particles to be ejected from the anterior pharynx rather than trapped at the end of the pump (Bhatla et al., 2015; Sando et al., 2021).

Particles trapped in the anterior isthmus are transported to the pharyngeal grinder via isthmus peristalsis, which occurs after a fraction of pumps (Song and Avery, 2012).

A number of ‘eat’ mutants with defective feeding behavior were identified by Avery (Avery, 1993a). Feeding phenotypes included slow pumping, defective timing of relaxation, and a “slippery pharynx” phenotype characterized by inefficient transport of particles. These feeding mutants were studied using video recordings at 60 frames per second (fps), a frame rate insufficient to resolve the rapid movements of the pharynx and food particles during muscle relaxation. As a result, aspects of pharyngeal behavior in feeding mutants have remained unclear. In particular, the mechanisms by which ‘slippery’ mutants are defective in anterior-to-posterior particle transport are poorly understood.

Here, we revisit the analysis of feeding mutants using high speed video microscopy at 1000 fps. We imaged worms feeding on bacteria and on polystyrene beads. By tracking the motions of pharyngeal components and particles within the pharyngeal lumen, we quantified the timing of contraction and relaxation, motion of particles in the pharyngeal lumen, and efficiency by which particles were transported and trapped. We also measured the pumping rate and the frequency of posterior isthmus peristalsis, by which particles trapped in the anterior isthmus are transported to the terminal bulb. Different slippery mutants showed a variety of behavioral defects, including weak isthmus contraction, failure to trap particles in the anterior isthmus, and abnormal timing of contractions and relaxations in different pharyngeal compartments. We also describe behavioral differences in slow-feeding and relaxation-defective mutants.

## Results

The class of slippery mutants were labeled as such by Avery because in these animals, bacteria were observed to slip toward the anterior instead of being trapped during pharyngeal relaxation (Avery, 1993a). We asked what differences in pharyngeal behavior relative to wild-type animals led to the slippery phenotype.

We first used our assay to describe wild-type pharyngeal behavior. In wild-type animals, pump timing and particle movement were consistent with previous findings (Fang-Yen et al., 2009). Each pump began with the onset of contraction of the corpus, followed 28.5 ± 14.8 ms (mean ± SD) later by onset of contraction of the anterior isthmus, followed 9.6 ± 11.7 ms (mean ± SD) later by onset of contraction of the terminal bulb (Fig, 1b, Video S1, Supplemental Table A3). Onset of relaxation occurred in the same order but faster, beginning with the corpus, followed 13.4 ± 9.9 ms later by the anterior isthmus, and then 6.4 ± 10.7 ms later by the terminal bulb (Fig. 1b). The pumping rate was 4.4 Hz ± 0.48 (mean ± SD). Isthmus peristalsis occurred after 22.6% of pumps, consistent with prior studies (Avery, 1993a; Song and Avery, 2012).

To assay the efficiency of particle transport, we reviewed video sequences to manually track trajectories of particles as they moved through the pharynx (see Methods). We found that pharynxes in wild-type animals transported particles efficiently: 79.5% (35 out of 44) of tracked particles in the corpus at the beginning of contraction reached the anterior isthmus and were trapped there at the end of the relaxation phase (Supplemental Table A4).

### Various behavioral defects contribute to inefficient transport in slippery mutants

We analyzed four slippery mutants: *eat-10, eat-13, eat-15*, and *eat-17*. By reviewing video recordings, we examined the patterns and timing of contraction and relaxation of pharyngeal compartments in each mutant. We also analyzed transport of particles (bacteria and/or polystyrene beads).

In wild-type animals, robust contraction of the anterior isthmus causes particles in the corpus to be drawn into the isthmus (Fig. 1, Fig. 2a,b) where they can be trapped. By contrast, in *eat-10* mutants the anterior isthmus frequently either failed to contract (18/49 pumps, 36%) or contracted weakly (14/49 pumps, 29%). Neither weak nor absent contractions were ever observed in wild-type worms. During weak contractions in *eat-10* worms, there was very limited (<20% of normal) opening of the pharyngeal lumen (Fig. 2c, Fig. 3, Video S2 worms 3-4). In cases of weak or absent anterior isthmus contraction, particles were not drawn into the anterior isthmus. As a result, in *eat-10* mutants, only 33% (16 out of 49) of the particles starting in the corpus at the beginning of a pump became trapped in the anterior isthmus at the end of the pump. In comparison, in wild-type animals, 79.5% moved from the corpus to isthmus at the end of the pump (Fig. 5). This finding is consistent with the original description of slippery pumping in which many particles reach the metacorpus at the peak of contraction but then return to the anterior procorpus during relaxation (Avery 1993a).

**Figure 2:**
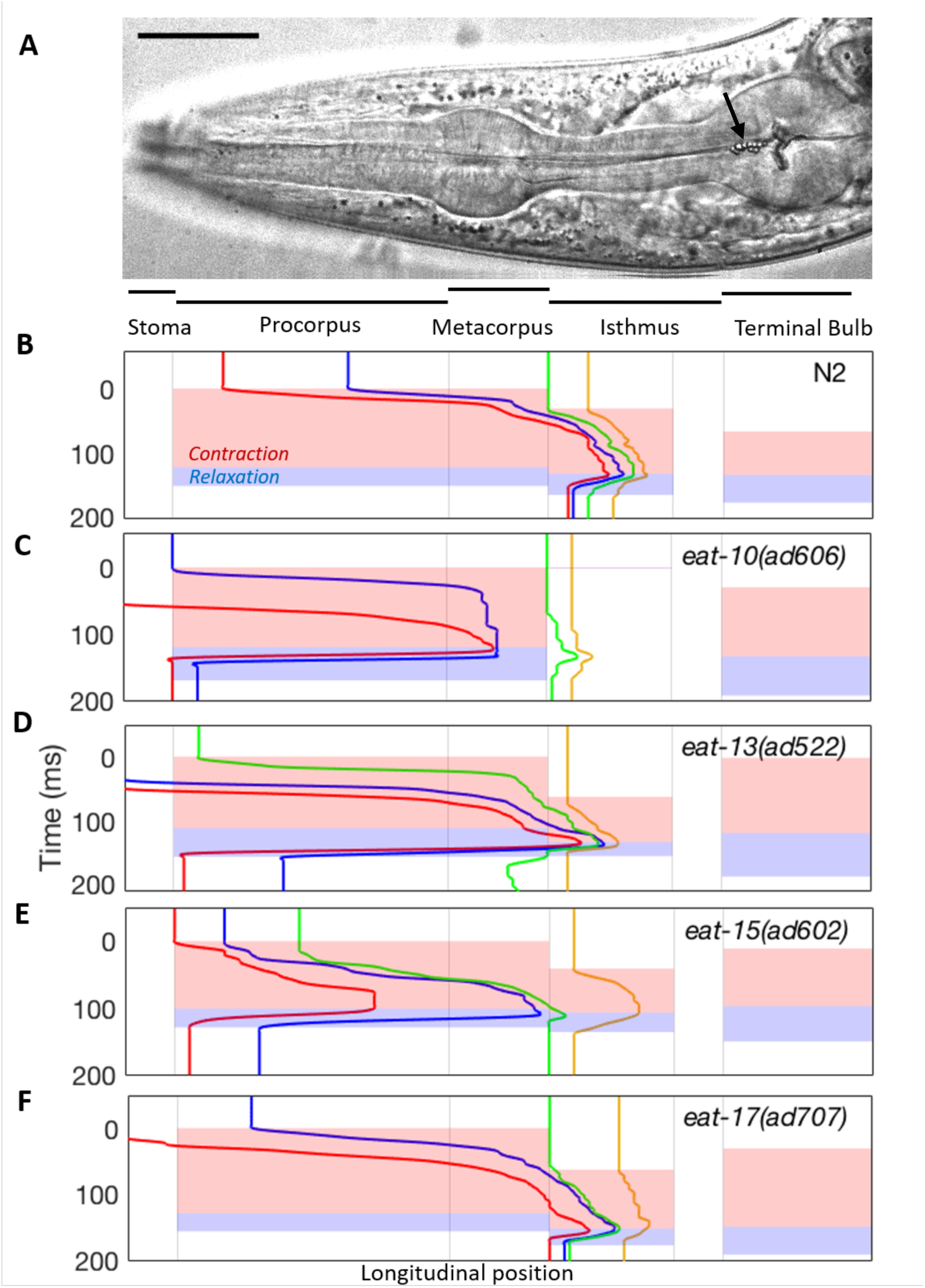
Particle tracking and pharyngeal subcomponent timing of wild type worms and slippery mutants. (A) The pharynx and pharyngeal compartments in a wild type animal. Scale bar: 25 µm. The particles just anterior to the TB, indicated by the arrow head, are polystyrene beads. (B-F) Pharyngeal muscle dynamics and particle tracking data from high-speed video microscopy of a single representative pump cycle from wild type (N2) and mutant worms. Colored traces indicate trajectories of four particles. Colored areas show timings of contraction and relaxation of the corpus, anterior isthmus, and terminal bulb, with red representing contraction and blue representing relaxation. The x axis corresponds to the longitudinal positions shown in (A). For eat-10, no isthmus contraction or relaxation was visible during this pump (Panel C, Supplemental Video S2 worm 4). Details of particle tracking data are given in Supplemental Table A4.

**Figure 3.**
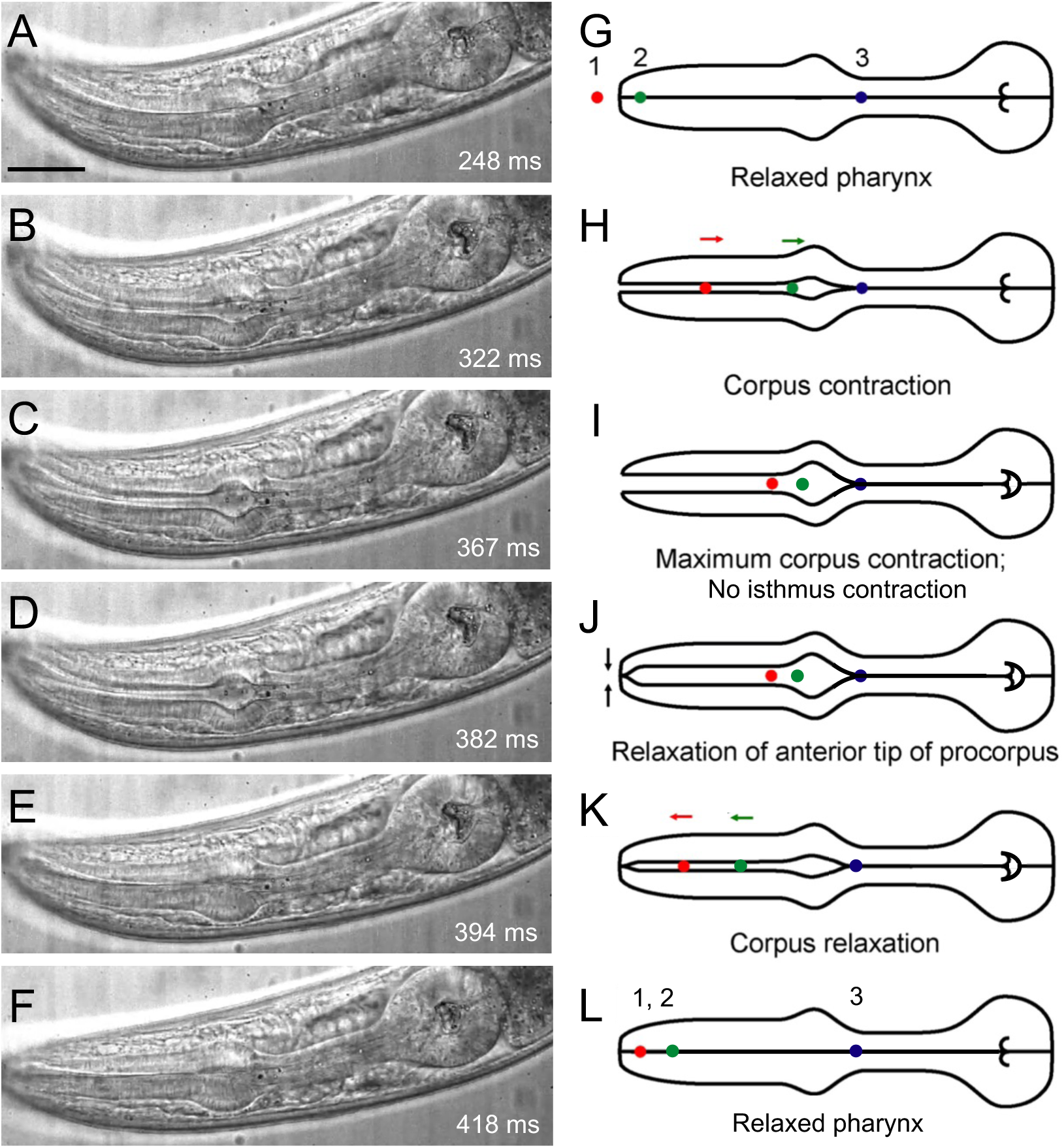
Particle transport in *eat-10* mutants. (A-F): Image sequence from high-speed video imaging of an *eat-10* worm (Video S2, worm 4). Scale bar: 25 µm. (G-L) Corresponding illustrations showing initial, intermediate, and final positions of three representative particles labeled 1 (red), 2 (green), and 3 (blue). During one cycle of contraction and relaxation, particle 1 has been transported from outside the pharynx to the anterior procorpus trap. Due to the lack of isthmus contraction, particle 2 has not progressed from the anterior procorpus trap to the anterior isthmus trap, but instead remains in the procorpus at the end of the pump cycle. Particle 3 remains in the anterior isthmus trap.

As in *eat-10* mutants, we found in *eat-15* worms a low (20/42 = 47.6%) rate of particle transport from the corpus to isthmus compared to *wild type* (Fig. 6). In some pumps, isthmus contraction was weak (Video S4 worm 3). Particle tracking showed that particles initially located in the corpus often did not enter the isthmus, instead stalling upon reaching the metacorpus (Fig. 4e). In some pumps, the particles moved back and forth along the lumen during relaxation, perhaps due to influence from movements in the terminal bulb (e.g. Video S4 worm 4).

**Figure 4:**
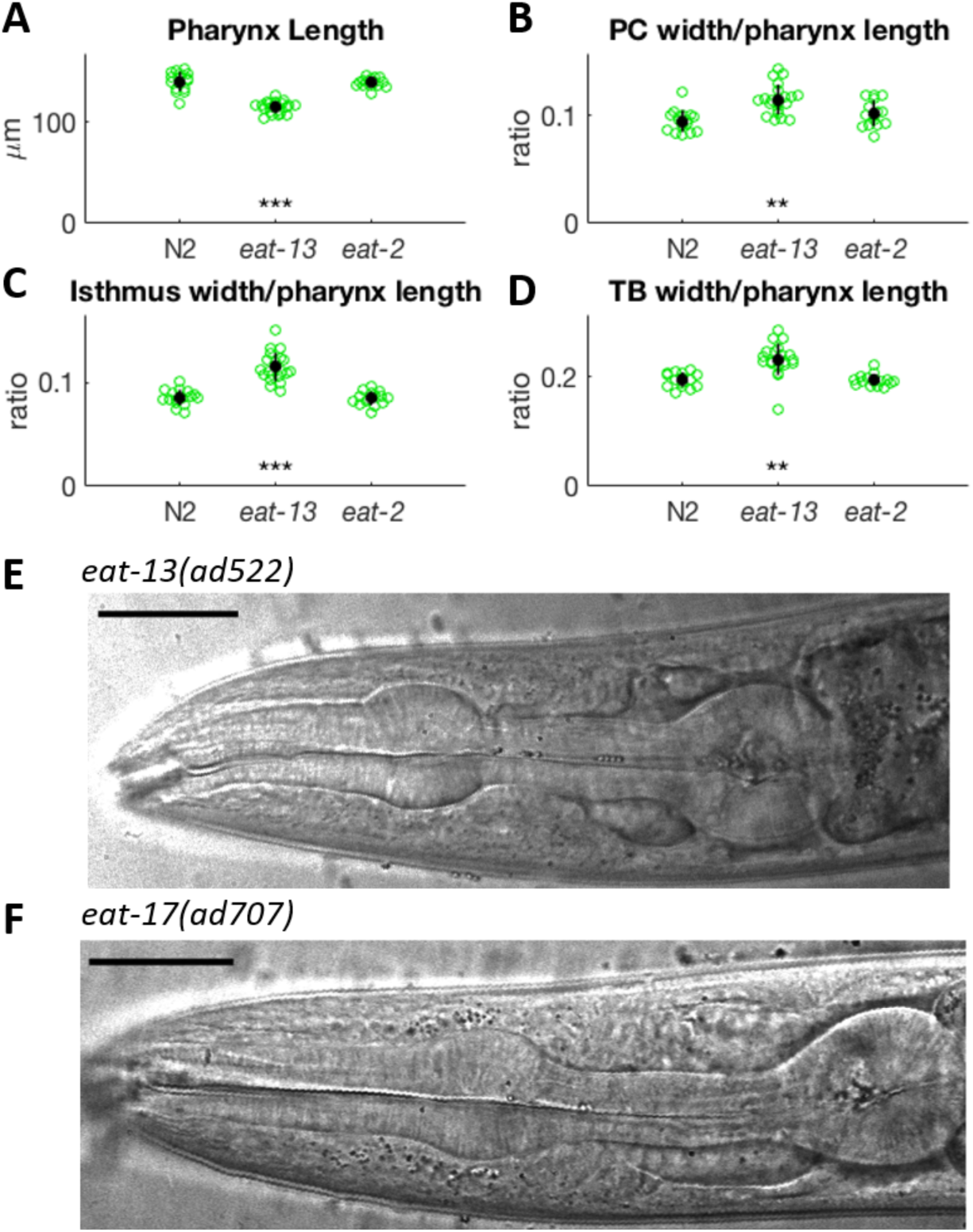
Pharynxes of *eat-13* animals are distorted. (A) Pharynx length for N2, *eat-13*, and *eat-2* animals. (B) Ratio of procorpus width to pharynx length. (C) Ratio of isthmus width to pharynx length. (D). Ratio of TB width to pharynx length. Significance threshold from t-test: * indicates p < 0.05, ** indicates p < 10^−5^, *** indicates p < 10^−8^. Note that the *eat-13* pharynx is generally shorter and wider than N2. N = 16 for N2, 21 for *eat-13*, and 15 for *eat-2*. See Supplemental Table A5 for detailed measurements. (E) Deformed *eat-13* pharynx. (F) Pharynx of an *eat-17* worm has a deformed grinder. See also Video S5. Scale bar: 25 µm for both images.

In *eat-13* mutants, efficiency of transport from the corpus to isthmus (24/38 = 63%) was only slightly lower than that of wild-type animals (79.5%). Pharynxes in *eat-13* mutants appeared physically distorted: the ratio of the isthmus width and the terminal bulb width to the length of the pharynx in *eat-13* mutants was greater than that ratio in wild-type worms (Video S3, Fig. 4a-e). This difference in morphology is unlikely to be explained by a nutritional defect alone since *eat-2* mutants, which have a defect in cholinergic excitation of the pharynx and therefore pump slowly (McKay et al., 2004; Raizen, Lee, and Avery, 1995), had a ratio no different from that of wild-type worms.

The gene *eat-17* encodes a GTPase-activating protein required for pharyngeal cuticle structure (Straud et al., 2013). In *eat-17* mutants, we observed a high percentage of particles failing to make posterior progress along the pharyngeal lumen (29% compared to 14% in wild-type, Supplemental Table A4). However, unlike *eat-13* mutants, *eat-17* mutants were highly efficient in corpus-to-isthmus transport, suggesting that most of the transport inefficiency in *eat-17* mutants was occurring in the isthmus. The grinder in the terminal bulb of *eat-17* mutant pharynxes appeared deformed (Video S5, Fig. 4f), as previously reported (Straud et al., 2013). Overall, *eat-17* mutants were able to transport particles from the corpus to the isthmus efficiently, but had defective transport within the isthmus (Video S5).

### Slippery mutants display altered isthmus and terminal bulb contraction timings

In wild-type worms, the onset of contraction of the terminal bulb nearly always occurred after onset of contraction in the isthmus, and the onset of relaxation of the terminal bulb occurred predominantly after onset of relaxation in the isthmus (Fig. 5, Supplemental Table A3). In *eat-10, eat-13*, and *eat-15* animals, the terminal bulb began contracting earlier on average than the isthmus (Fig. 5a). Similarly, on average, the terminal bulb also began relaxation before the isthmus (Fig. 5b), although the difference was smaller than that of the contraction delay.

**Figure 5:**
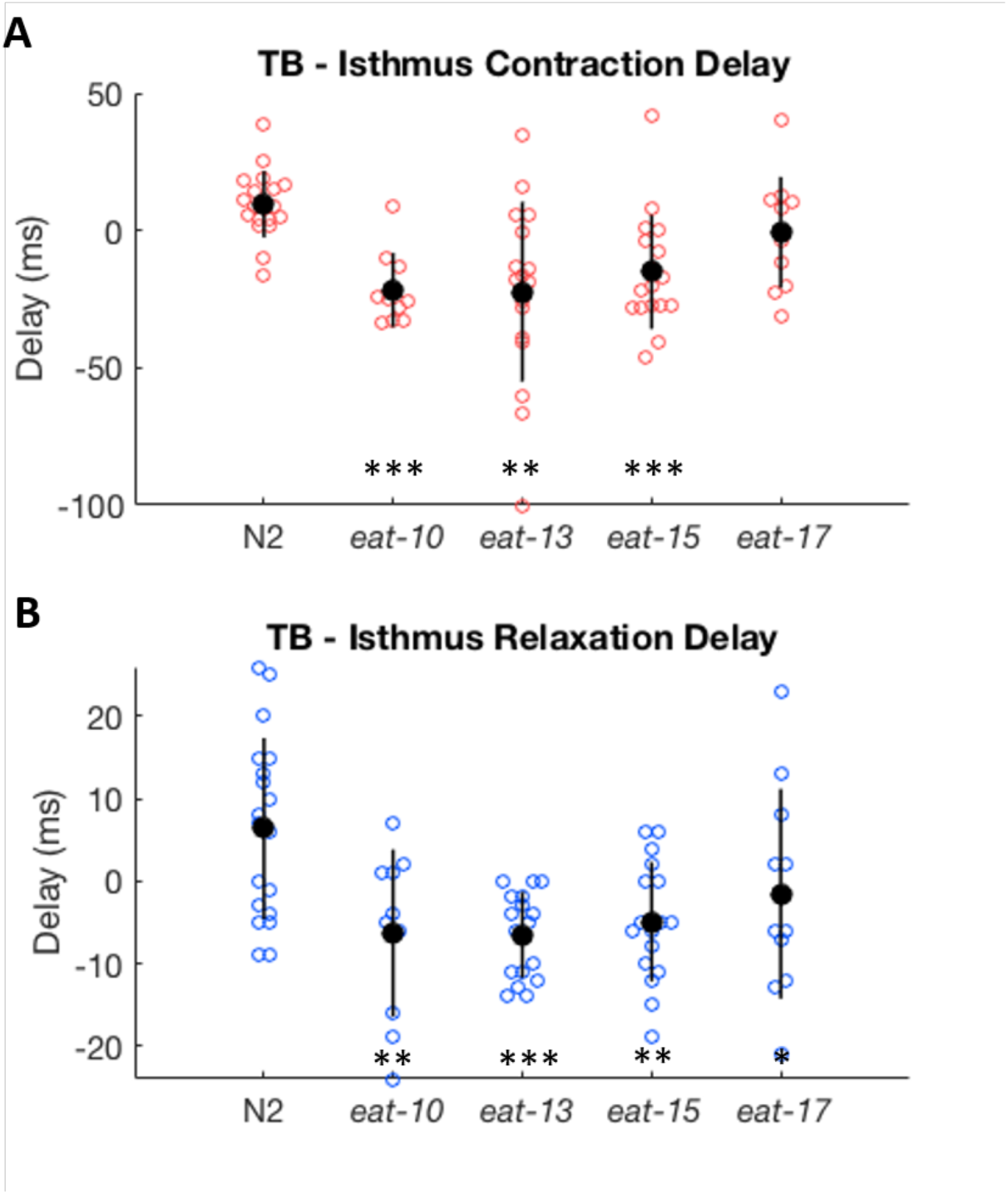
Delay between the onset of (A) contraction and (B) relaxation of the terminal blub (TB) relative to the isthmus in wild-type worms and slippery mutants. Negative delay indicates that terminal bulb contracted/relaxed prior to isthmus. Each point represents a single pump. For each genotype, data is from 1 pump from each of N = 10-19 animals (See Supplemental Table A3). Error bars indicate +/- SEM. Results of two-tailed Student’s t-test comparing each strain to N2; * indicates p<0.05, ** indicates p<0.005, *** indicates p<0.0005.

### Relaxation defective mutants show defective transport

The pharynxes of loss-of-function mutants e*at-6* fail to relax completely during the pump cycle (Avery, 1993a). The gene *eat-6* encodes a sodium/potassium ATPase, a pump required to maintain the resting membrane potential (Davis et al., 1995). In *eat-6* mutants, we observed that the terminal bulb only occasionally came back to its fully relaxed position, usually appearing to only partially relax (Video S6). In addition to this failure of the terminal bulb to fully relax, the relaxation phases were proportionally longer in *eat-6* mutants than in the *wild type*. Corpus (p=0.001) and isthmus (p=0.02) contraction phases were significantly shorter in *eat-6* after normalizing for pump period, indicating that relaxation made up a greater percentage of the overall pump. Like *eat-10* animals, *eat-6* worms frequently showed weak isthmus contractions, precluding effective transport (Fig. 7c, Video S6 worms 2-4). This resulted in a lower rate of corpus to isthmus transport (23.2% compared to 79.5% in N2; Fig. 6).

**Figure 6:**
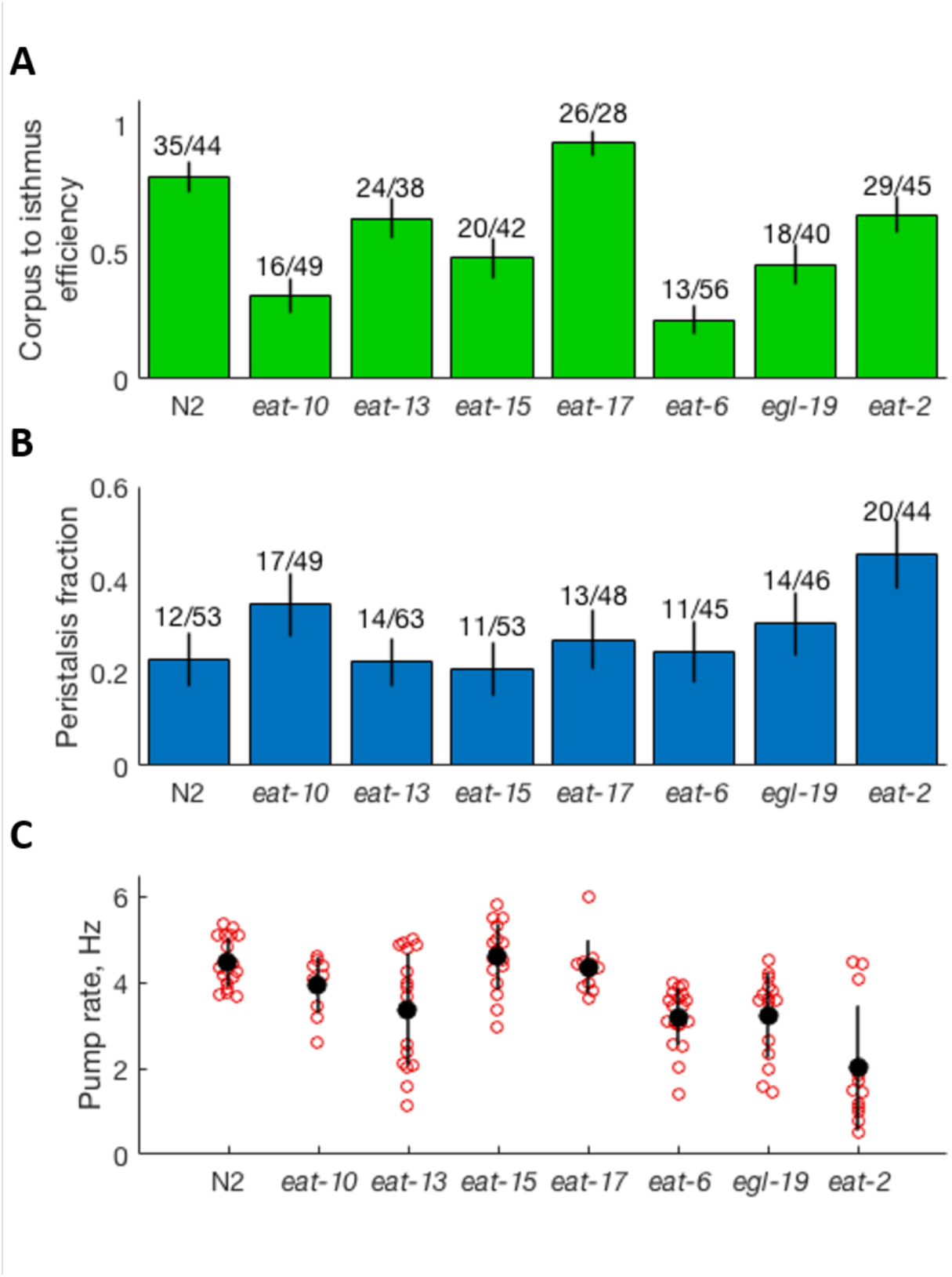
(A) Particle transport efficiency, defined as the fraction of particles starting in the corpus that are transported to the isthmus during a pump. Error bars show ± standard deviation assuming a binomial distribution. (B) Fraction of pumps that were followed by isthmus peristalsis. Error bars show standard deviation assuming a binomial distribution. (C) Pump rate, with black bars indicating mean +/- SD; n = 11-20. Additional data can be found in Supplemental Tables A1 and A4.

**Figure 7.**
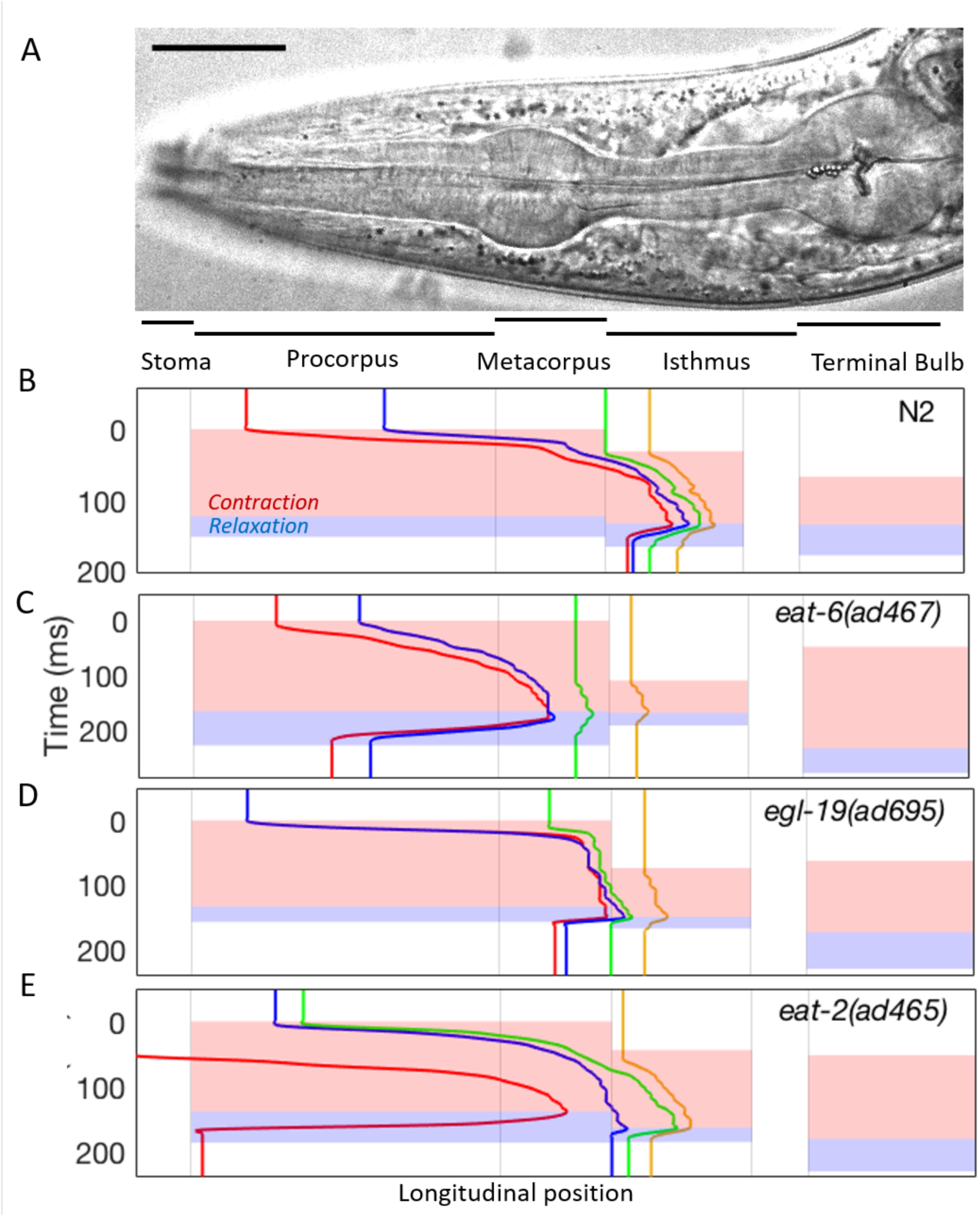
Particle tracking and pharyngeal subcomponent timing of N2, eat-6, egl-19, and eat-2 mutants. (A-E) Representative particle trajectories and contraction/relaxation timings, as in Fig. 2.

Like *eat-6* mutants, gain-of-function mutants of *egl-19 (*an L-type voltage-gated calcium channel (Lee et al 1997) fail to relax completely during the pump cycle (Avery, 1993a). In an *egl-19* gain-of-function mutant, the most notable behavior was occasional (3/46 pumps) isthmus and terminal bulb “yawns” in which those parts of the pharynx would remain contracted while the corpus went through its normal pump cycle (Video S7 worm 1-2). These terminal bulb contractions would span two pumps of the corpus. Even when the contraction ended prior to the next pump (i.e., no discernable “yawn” occurred), the isthmus and terminal bulb remained contracted longer than observed in wild-type worms (Supplemental Table A2). Consistent with this, we found a longer delay between onset of corpus relaxation, and that of the isthmus, and subsequently the terminal bulb (Supplemental Table A3). We observed a lower rate of corpus to isthmus transport (18/40, 45%), as well as an increase in particles ending in the metacorpus (21% compared to 0% in wild type (Fig. 6, Supplemental Table A4).

### Slow pumping mutant *eat-2* shows normal pharyngeal dynamics and particle transport

We next analyzed animals mutant for the gene *eat-2*, which encodes a non-alpha subunit of a nicotinic acetylcholine receptor (McKay et al., 2004). Because EAT-2 is required for excitation of the pharynx by the motor neuron MC (Raizen et al., 1995) animals mutant for *eat-2* have a pump rate much lower than that of wild-type (1.3 ± 0.5 Hz compared to 4.4 ± 0.48 Hz for N2). To assess differences in pump timing, we normalized timing measurements by pump duration. After normalizing for pump duration, we found that the duration of the contraction and relaxation phases were nearly identical to that of N2. Consistent with previously reported results (Kozlova et al., 2019), we observed a higher rate of peristalsis in *eat-2*, with peristalsis occurring after 45.5% of pumps, compared to 22.6% in N2. Aside from slow pumping and increased peristalsis, *eat-2* mutants displayed normal pharyngeal muscle dynamics and particle trapping (Fig. **7E**, Video S8). Transport efficiency was similar to that of wild-type worms, with about 65% of particles in the corpus becoming trapped in the isthmus.

## Discussion

In this work we used high speed video microscopy to observe pharyngeal muscle movements and particle transport in feeding mutants. We found that three of the four mutants described as slippery had defects in posteriorward transport, consistent with previous observations (Avery, 1993a).

The mechanisms by which differences in pharyngeal movements create particle transport defects were clear in only a few cases. In *eat-10*, and possibly *eat-15*, inefficiency of posterior transport could be explained by a deficiency or absence of anterior isthmus contraction. Isthmus contraction is required for particles to be drawn from the metacorpus into the isthmus, prior to being trapped there.

Though pharyngeal muscles are electrically coupled (Starich et al., 1996) the pharynx exhibits highly compartmentalized muscle dynamics, between and even within individual muscle cells. For example, the anterior tips of the pm3 muscle cells, which span the procorpus, relax earlier than the rest of the corpus during pumping (Fang-Yen et al., 2009), a behavior that is antagonized by activity of the M1 neuron (Bhatla et al., 2015; Sando et al., 2021). Similarly, although the isthmus is composed of just one muscle cell type (pm5), which spans the length of the isthmus, the anterior and posterior compartments of the isthmus display different behaviors: the anterior isthmus contracts during pumps whereas the posterior isthmus displays a peristaltic movement after every fourth or five pumps. The finding of weak isthmus pumping but normal posterior isthmus peristalsis in *eat-10* (and possibly *eat-15* mutants) suggests that muscle contractions in subcellular compartments of the pharyngeal isthmus are mediated by distinct mechanisms.

In discussing reasons for isthmus peristalsis, Avery proposed that normal corpus and terminal bulb function require them to be hydrodynamically isolated from each other during pumping (Avery, 1993c). During corpus contraction, pharyngeal lumen pressure decreases, and the bacteria suspension is drawn into the pharyngeal lumen from the stoma. The posterior isthmus remaining closed (relaxed) at this time may serve to reduce gastro-pharyngeal reflux, i.e. fluids and/or particles drawn into the pharynx from the intestine. However, if the relaxed position of the posterior isthmus mechanically isolates the terminal bulb from the corpus, then the timing of terminal bulb contraction should not affect corpus to anterior isthmus transport. In all four slippery mutants studied, we observed a premature terminal bulb relaxation (Fig. 5). This observation suggests that there is a connection between terminal bulb contraction and particle transport efficiency, and that the TB is not fully isolated from the corpus and anterior isthmus. It is possible that a delay between terminal bulb and corpus contraction reduces the fluid resistance of the posterior pharynx, and so desynchronization of the terminal bulb contraction relative to that of the corpus and isthmus normally helps to maintain posteriorward flow.

We observed poor pharyngeal transport in animals mutant for *eat-17*, which encodes a GTPase-activating protein required for pharyngeal cuticle structure (Straud et al., 2013). The observation of poor particle transport despite no detectable defect in pharyngeal motion timing suggests that the pharyngeal cuticle may play a role in transport, perhaps via mechanical interactions with food particles. Alternatively, poorly ground bacteria in the terminal bulb due to *eat-17* grinder dysfunction may affect the mechanics of bacterial transport in the entire pharynx, again perhaps because the terminal bulb is not fully mechanically isolated from the corpus and anterior isthmus.

We found that *eat-6* and *egl-19* mutants, previously described as defective in relaxation (Avery 1993a), were most notable for their extended contraction phases. We also found that both of these mutants were deficient in posteriorward particle transport. Therefore, these relaxation-defective mutants can also be described as slippery. Considering the importance of the relaxation phase for trapping and transport, this result is not surprising.

In this work we have focused on describing the behavioral defects in feeding mutants. Future efforts will aim to connect the molecular identity of specific genes to specific behavioral defects. Computational modeling of pressure gradients and fluid flow within the pharyngeal lumen may help connect behavioral defects to particle transport, particularly for timing defects. The future use of genetically encoded calcium or voltage sensors could allow for an assessment of excitation in different muscle compartments. The combination of genetic analysis and high-resolution phenotyping of pharyngeal behavior (for example, of *eat-10*), as well as optogenetic manipulation of neural activity (Trojanowski et al., 2016, 2014) could lead to new insights into excitable cells and organs, in particular regarding the mechanisms of subcellular compartmentalization of muscle contraction.

## Material and Methods

Strains used in this study are shown in Figures 2 and 7. *C. elegans* were cultured using standard methods (Stiernagle, 2006). For experiments, 5-10 young adult hermaphrodite animals were transferred to a 1 mm thick pad composed of 2% agarose in NGM buffer on a microscope slide, in addition to 1-2 µL of either an OP50 *E. coli* suspension in LB medium or a 1% (vol/vol) suspension of 1 µm diameter polystyrene beads (Polysciences 431), diluted in NGM buffer to 0.1% vol/vol. NGM buffer consisted of the same components as NGM media (Stiernagle, 2006) without agar, peptone, or cholesterol. The pad was covered with a #1 cover slip and worms were given 15-45 minutes to acclimate to the pad before imaging. We mounted slides on an inverted microscope (Nikon TE-2000S) with a 60X Plan Apo (NA 1.4) oil immersion objective lens under differential interference contrast optics. For each animal we saved 1-3 recordings, each 4-5 s in duration, at 1000 frames per second (fps) using a Vision Research Phantom V9.1 high speed camera.

To record muscle timing data, we used a custom MATLAB script to review videos and manually recorded the frame numbers corresponding to 10 events: the start of corpus contraction, the start of corpus relaxation, the end of corpus relaxation, the same for both the isthmus and the terminal bulb, and the beginning of the next pump. This was done only for recording periods in which all parts of the pump cycle were clearly visible. We manually scored each pump for whether or not it was followed by an isthmus peristalsis.

To perform particle tracking, we used the same MATLAB program to analyze videos frame by frame and recorded the approximate locations of several particles in each pump, as well as the timings of each particle’s beginning, peak, and end of movement. To approximate particle locations, we established several guideposts along the pharynx (tip of the corpus, both ends of the metacorpus, the isthmus trap, the end of the anterior isthmus, and the grinder) and assigned them approximate distances relative to the tip of the stoma. We tracked at least one particle in the corpus and isthmus on every pump in which there were particles in those compartments.

All statistical analyses were performed in Python. Because duration of components of the pharyngeal pump is affected by the total pump time (Fang-Yen et al., 2009), we normalized time measurements by the pump period for each trial before making statistical comparisons between strains. To assess the significance of these particle tracking samples, we used the 2×2 Fisher test, comparing each mutant strain with N2. All pump timing data can be found in Supplemental Tables A1-3, and all particle tracking data can be found in Supplemental Table A4.

## Acknowledgements

Some strains were provided by the *Caenorhabditis* Genetics Center (CGC), which is funded by NIH Office of Research Infrastructure Programs (P40 OD010440). We thank Levi Kanu for conducting preliminary experiments in this project.

## Videos

https://doi.org/10.6084/m9.figshare.16713268

**Video S1. High speed video of pharyngeal pumping in N2 animals**

Pump cycles from 4 wild type worms feeding on 1 µm beads. The beginning of each animal’s recording can be found at the following time points (m:s): Worm 1: 0:00; Worm 2: 0:13; Worm 3: 0:27; Worm 4: 0:42. Scale bar = 25 µm in all videos. Worm 1 corresponds to tracking plot in Fig. 2b. Four particles are annotated during the first two pumps to highlight their movements.

https://doi.org/10.6084/m9.figshare.16713265

**Video S2. High speed video of pharyngeal pumping in *eat-10* animals**

The beginning of each animal’s recording can be found at the following time points (m:s): Worm 1: 0:00; Worm 2: 0:15; Worm 3: 0:30; Worm 4: 0:45. Note the weak isthmus contractions in worms 3 and 4. Worm 4 corresponds to tracking plot in Fig. 2c.

https://doi.org/10.6084/m9.figshare.16732798

**Video S3. High speed video of pharyngeal pumping in *eat-13* animals**

The beginning of each animal’s recording can be found at the following time points (m:s): Worm 1: 0:00; Worm 2: 0:12; Worm 3: 0:28; Worm 4: 0:41. Note the deformed pharynx in these worms. Worm 2 corresponds to tracking plot in Fig. 2d.

http://doi.org/10.6084/m9.figshare.16713253

**Video S4. High speed video of pharyngeal pumping in *eat-15* animals**

The beginning of each animal’s recording can be found at the following time points (m:s): Worm 1: 0:00; Worm 2: 0:16; Worm 3: 0:29; Worm 4: 0:40. Worm 3 has weak isthmus contraction. Worm 4 shows particles moving back and forth along the lumen during relaxation, perhaps due to the influence of the terminal bulb. Worm 2 corresponds to tracking plot in Fig. 2e.

https://doi.org/10.6084/m9.figshare.16713262

**Video S5. High speed video of pharyngeal pumping in *eat-17* animals**

The beginning of each animal’s recording can be found at the following time points (m:s): Worm 1: 0:00; Worm 2: 0:12; Worm 3: 0:27; Worm 4: 0:44. Note the deformed pharynxes in these worms. The grinder flaps are hard to resolve, indicating a potential defect in grinder formation as reported (Straud et al., 2013). Worm 3 corresponds to tracking plot in Fig. 2f.

https://doi.org/10.6084/m9.figshare.16713259

**Video S6. High speed video of pharyngeal pumping in *eat-6* animals**

The beginning of each animal’s recording can be found at the following time points (m:s): Worm 1: 0:00; Worm 2: 0:20; Worm 3: 0:48; Worm 4: 1:12. Note that the terminal bulb does not come back to the fully relaxed position in these worms, but remains slightly contracted even when the other compartments have closed. Worms 2-4 also have weak isthmus contraction. Worm 2 corresponds to tracking plot in Fig. **7c**.

https://doi.org/10.6084/m9.figshare.16713274

**Video S7. High speed video of pharyngeal pumping in *egl-19* animals**

The beginning of each animal’s recording can be found at the following time points (m:s): Worm 1: 0:00; Worm 2: 0:16; Worm 3: 0:40; Worm 4: 0:58. Note the unique “yawns” of the terminal bulb in worms 1 and 2. In those pumps, the terminal bulb remained fully contracted for two pump cycles. Worm 3 corresponds to tracking plot in Fig. 7d.

https://doi.org/10.6084/m9.figshare.16713271

**Video S8. High speed video of pharyngeal pumping in *eat-2* animals**

The beginning of each animal’s recording can be found at the following time points (m:s): Worm 1: 0:00; Worm 2: 0:27; Worm 3: 0:47; Worm 4: 1:15. Note the extended delay between pump cycles in these worms compared to N2. Worm 2 corresponds to tracking plot in Fig. 7e.

## Supplemental Tables

https://doi.org/10.6084/m9.figshare.16733707

**Supplemental Table A1.** Pump timing summary data.

| Strain | N | Duration (STD), ms | Period (STD), ms | Frequency (STD), ms |
| --- | --- | --- | --- | --- |
| *N2* | 19 | 181.7 (24.6) | 227.2 (27.7) | 4.4 (0.48) |
| *eat-10* | 10 | 199.6 (25.3) | 262.1 (51.0) | 3.82 (0.63) |
| *eat-13* | 17 | 191.8 (26.7) | 358.1 (179.7) | 2.79 (0.93) |
| *eat-15* | 17 | 165.6 (23.4) | 224.5 (42.5) | 4.45 (0.7) |
| *eat-17* | 11 | 180.3 (24.1) | 233.3 (27.3) | 4.29 (0.45) |
| *eat-6* | 20 | 229.0 (38.9) | 331.9 (103.4) | 3.01 (0.71) |
| *egl-19* | 15 | 248.3 (54.5) | 350.5 (145.2) | 2.85 (0.83) |
| *eat-2* | 12 | 222.4 (26.6) | 766.6 (485.1) | 1.3 (0.5) |

**Supplemental Table A2.** Pump phase lengths. CC: Corpus contraction, CR: Corpus relaxation, IC: Isthmus contraction, IR: Isthmus relaxation, TC: Terminal Bulb contraction, TR: Terminal Bulb relaxation.

| Strain | CC (STD), ms | CR (STD), ms | IC (STD), ms | IR (STD), ms | TC (STD), ms | TR (STD), ms |
| --- | --- | --- | --- | --- | --- | --- |
| *N2* | 109.7 (19.6) | 34.8 (11.0) | 94.6 (18.1) | 34.9 (15.2) | 91.4 (13.6) | 52.2 (14.5) |
| *eat-10* | 121.1 (15.2) | 44.4 (15.2) | 100.2 (12.6) | 43.1 (14.1) | 115.6 (20.2) | 63.0 (17.3) |
| *eat-13* | 108.8 (21.3) | 35.6 (8.7) | 98.0 (14.0) | 35.3 (11.0) | 113.8 (32.4) | 61.7 (16.3) |
| *eat-15* | 110.2 (21.2) | 30.5 (7.8) | 85.6 (19.7) | 31.7 (6.6) | 95.8 (17.4) | 47.2 (7.6) |
| *eat-17* | 110.9 (15.0) | 32.3 (5.1) | 106.8 (16.8) | 35.9 (11.4) | 106.0 (28.5) | 49.8 (16.6) |
| *eat-6* | 124.0 (24.0) | 52.6 (17.6) | 109.7 (22.2) | 53.1 (18.1) | 116.2 (27.5) | 83.2 (28.5) |
| *egl-19* | 132.3 (25.3) | 38.7 (17.4) | 130.5 (37.0) | 38.1 (11.5) | 143.8 (33.1) | 69.5 (18.7) |
| *eat-2* | 125.0 (29.9) | 46.7 (9.9) | 110.8 (26.9) | 40.8 (7.9) | 117.6 (28.0) | 56.6 (12.0) |

**Supplemental Table A3.**
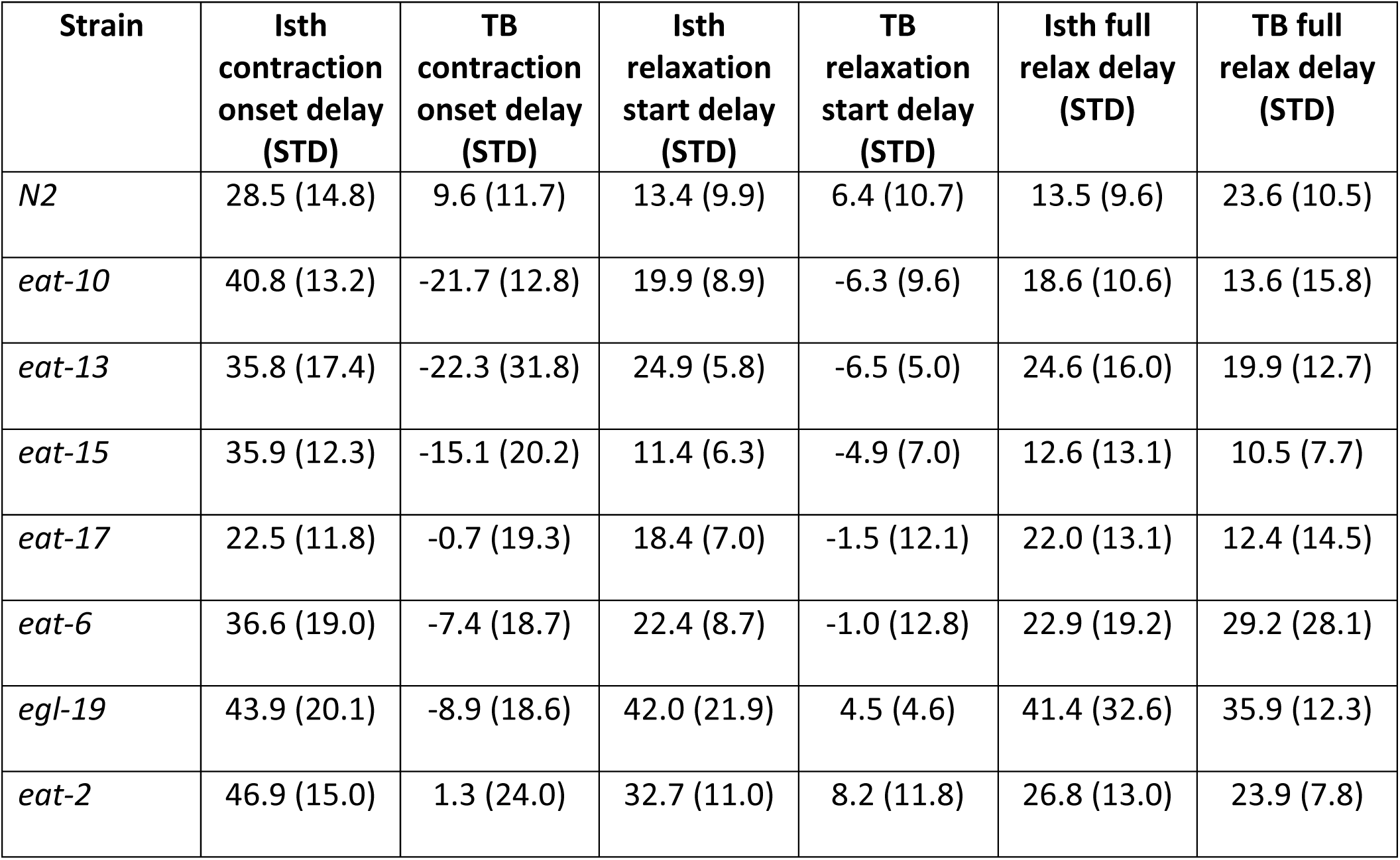
Isthmus and Terminal bulb (TB) delays. Isthmus delays are relative to the corpus behavior, and terminal bulb delays are relative to isthmus behavior, with negative values indicating that isthmus behavior occurred before corpus, or TB behavior before isthmus. All data are in ms.

| Strain | Isth contraction onset delay (STD) | TB contraction onset delay (STD) | Isth relaxation start delay (STD) | TB relaxation start delay (STD) | Isth full relax delay (STD) | TB full relax delay (STD) |
| --- | --- | --- | --- | --- | --- | --- |
| *N2* | 28.5 (14.8) | 9.6 (11.7) | 13.4 (9.9) | 6.4 (10.7) | 13.5 (9.6) | 23.6 (10.5) |
| *eat-10* | 40.8 (13.2) | -21.7 (12.8) | 19.9 (8.9) | -6.3 (9.6) | 18.6 (10.6) | 13.6 (15.8) |
| *eat-13* | 35.8 (17.4) | -22.3 (31.8) | 24.9 (5.8) | -6.5 (5.0) | 24.6 (16.0) | 19.9 (12.7) |
| *eat-15* | 35.9 (12.3) | -15.1 (20.2) | 11.4 (6.3) | -4.9 (7.0) | 12.6 (13.1) | 10.5 (7.7) |
| *eat-17* | 22.5 (11.8) | -0.7 (19.3) | 18.4 (7.0) | -1.5 (12.1) | 22.0 (13.1) | 12.4 (14.5) |
| *eat-6* | 36.6 (19.0) | -7.4 (18.7) | 22.4 (8.7) | -1.0 (12.8) | 22.9 (19.2) | 29.2 (28.1) |
| *egl-19* | 43.9 (20.1) | -8.9 (18.6) | 42.0 (21.9) | 4.5 (4.6) | 41.4 (32.6) | 35.9 (12.3) |
| *eat-2* | 46.9 (15.0) | 1.3 (24.0) | 32.7 (11.0) | 8.2 (11.8) | 26.8 (13.0) | 23.9 (7.8) |

**Supplemental Table A4.**
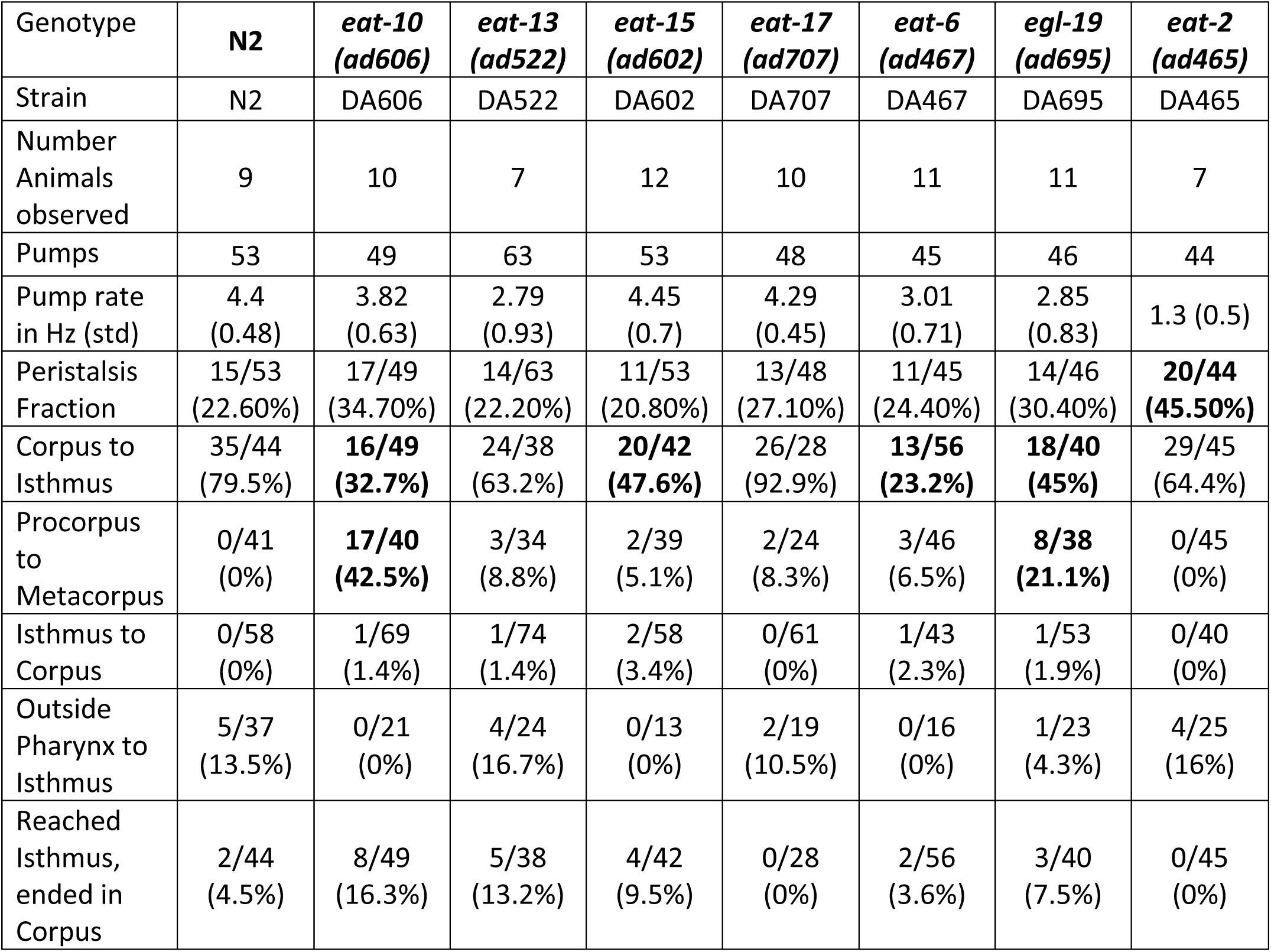
Results of particle tracking analysis. Bold cells indicates result of 2×2 Fisher test for each strain compared to N2 with p < 0.05.

**Supplemental Table A5.** Selected pharynx measurements for N2, *eat-13*, and *eat-2*. All cells show mean of measurements +/-- SD. *eat-2* was assayed to account for nutritional deficiencies that may arise from inefficient feeding. Results of two-tailed t-test given, with * = p < 0.05, ** = p < 10^−5^, *** = p < 10^−8^

| Strain | n | Total pharyngeal length (μm) | Procorpus diameter (μm) | Metacorpus diameter (μm) | Isthmus diameter (μm) | Terminal bulb diameter (μm) |
| --- | --- | --- | --- | --- | --- | --- |
| N2 | 16 | 140.675 ± 9.607 | 13.379 ± 1.541 | 22.393 ± 2.427 | 12.092 ± 1.376 | 27.437 ± 2.887 |
| *eat-13* | 21 | 114.862 ± 6.012*** | 13.212 ± 1.573 | 21.697 ± 1.856 | 13.219 ± 1.543 | 26.611 ± 3.399 |
| *eat-2* | 15 | 139.458 ± 4.684 | 14.246 ± 1.438 | 23.476 ± 1.499 | 11.849 ± 0.946 | 26.971 ± 1.393 |
|  |  |  | Procorpus Diam/Pharynx length | Metacorpus Diam/Pharynx Length | Isthmus diam/Pharynx Length | Terminal bulb Diam/Pharynx Length |
| N2 |  |  | 0.095 ± 0.01 | 0.159 ± 0.01 | 0.086 ± 0.008 | 0.195 ± 0.013 |
| *eat-13* |  |  | 0.115 ± 0.014** | 0.189 ± 0.014*** | 0.115 ± 0.014*** | 0.232 ± 0.029** |
| *eat-2* |  |  | 0.102 ± 0.012 | 0.168 ± 0.009* | 0.085 ± 0.007 | 0.194 ± 0.011 |

